# When Attended and Conscious Perception Deactivates Fronto-Parietal Regions

**DOI:** 10.1101/097410

**Authors:** Ausaf Ahmed Farooqui, Tom Manly

## Abstract

Conscious and attended perception is commonly thought to elicit fronto-parietal activity. However, supportive evidence comes largely from studies which involve detecting a target and reporting its visibility. This approach confounds conscious perception with goal completion of either the perceptual task of detection or the metacognitive task of introspective reporting. In contrast, in real life such perceptions are a means of achieving goals and rarely a goal in themselves, and almost never involve explicit metacognitive reports. It therefore remains unclear if fronto-parietal activity is indeed a correlate of conscious perception or is the result of confounds related to goal completion. Here we show that conscious and attended perception when delinked from goals does not increase fronto-parietal activity, and when inconsequential for the goal may even deactivate these regions. In experiments 1 and 2 participants attended to a highly visible stream of letters to detect the occasional targets in their midst. The non-target letters, in spite of being visible and attended to, deactivated fronto-parietal regions. In experiment 3 we looked at the activity elicited by a loud auditory cue that had to be kept in memory for up to 9 s and used to select the correct rule for completing the goal. Even such a salient, attended and remembered event did not elicit prefrontal activity. Across these experiments conscious and attended perception only activated the relevant sensory regions while goal completion events activated fronto-parietal regions.

**Significance statement:** Consciousness and attended perception has been seen to correlate with fronto-parietal activity. This informs key theories of consciousness and attention, e.g. widespread availability of incoming information or its higher level representation causes perceptual awareness, or that top down attention during perception broadcasts incoming sensations into frontal and parietal regions. However such experiments unwittingly conflate attended and conscious perception with some form of goal completion, whereas such perception in our daily life mostly serves as a means of goal completion and not a goal in itself. Here we show that such perception when delinked from goal completion does not activate fronto-parietal regions, and may even deactivate these regions if the percept is inconsequential for goal completion.

## Introduction

Certain perceptions have a special status. Not only do they induce activity in our neural systems and influence our behavior, but we have a subjective experience of perceiving them and we know that we have perceived them. Delineating the neural correlates of conscious perception (NCC) is considered a tractable first step in understanding how and why some stimuli reach conscious awareness (Crick, 1995). A key debate in this concerns whether widespread fronto-parietal activity is necessary for being aware of the incoming percept (e.g. Dehaene, 2014 and Koch et al. 2016).

A number of studies have linked consciousness of the percept to greater and more widespread fronto-parietal activity (for reviews see Rees, 2007; Dehaene and Changeux, 2011), and form a key basis of both neural and cognitive theories of consciousness. For example, consciousness is thought to arise through higher level cognitions that re-represent the first order perceptual state, or alternately, through a global neuro-cognitive broadcast of local perceptual information (Lau and Rosenthal, 2011; Baars, 2013). However, the vast majority of these empirical findings come from experiments where conscious perception ends up being the *goal* of the task being executed, making it unclear if conscious awareness and not goal completion was the actual elicitor of the fronto-parietal activity.

In masking experiments, for example, where subjects have to explicitly detect degraded and fleeting stimuli, the conscious perception of the otherwise difficult to perceive event is the obvious goal of the task subjects pursue (e.g. Lau and Passingham, 2007). Likewise, when participants monitor and report changes in their percept during binocular rivalry, detecting and reporting the change becomes the goal of the task (Lumer, Friston & Rees, 1998; Knapen et al. 2011). Furthermore, the requirement to report awareness creates an additional metacognitive task that requires an explicit introspective monitoring of the ongoing phenomenal state. Conscious perception in these designs will be linked to success in difficult perceptual inference, goal attainment and metacognitive access (see also Aru et al. 2012; de Graaf et al. 2012). Each of these is known to involve fronto-parietal activity (Summerfield and Egner, 2009; Crittenden et al. 2013; Ericsson et al. 2008; Farooqui et al. 2012; Fleming and Dolan, 2012).

Relatedly, studies on attention typically focus on the goals of visual search, i.e. targets, and show that they elicit strong fronto-parietal activity (Corbetta & Shulman, 2002; Dehaene et al. 2006; Duncan, 2006). But top down attention is never limited to goals, all task relevant events (including objects and spaces) that are instrumental in goal completion, unlike events unrelated to the task, are attended albeit to a lesser extent than goals. Hence, when we search through a pile of papers, not just the target paper but the entire pile being searched through, has to be sequentially attended to, unlike other parts of the desk. The fate of attended task events that are not goals is unclear.

Across three experiments we investigate the neural fate of perceptual events that were easy to perceived, unambiguously conscious, had to be attended to as part of task execution but were not goals in themselves. We find that such events only activated their specific sensory regions, and could even deactivate fronto-parietal regions.

## Methods

In the first two experiments we created an experimental equivalent of searching for a specific paper in a pile of papers. Participants attentively searched through serial letter presentations for pre-specified targets in their midst. We investigated the nature of activity elicited by individual non-target letter events that were highly visible and had to be attended to to detect targets in their midst. In experiment 1 we looked at the effect of top-down attention on the activity elicited by these letters, while in experiment 2 we compared activity during periods of such attentive and conscious perception to that during periods of passive wait in front of a blank screen (experiment 2).

Participants saw sequences of highly visible letters presented at the rate of 1/s (on for 900 ms) for the occurrence of the occasional prespecified letter targets interspersed with, and of the same size, color and font as, these letters (Figure 1). A three-letter word (e.g., ‘CAT’; on for 3.5 s) presented at the start of the trial indicated the three targets (T1, T2 T3; e.g., Fig. 1a, T1 ‘D’, T2 ‘A’, T3 ‘T’). The letter sequence began after a jittered gap of 1—5 s. The letter stream consisted of a total of 40 letter presentations.

**Figure 1.**
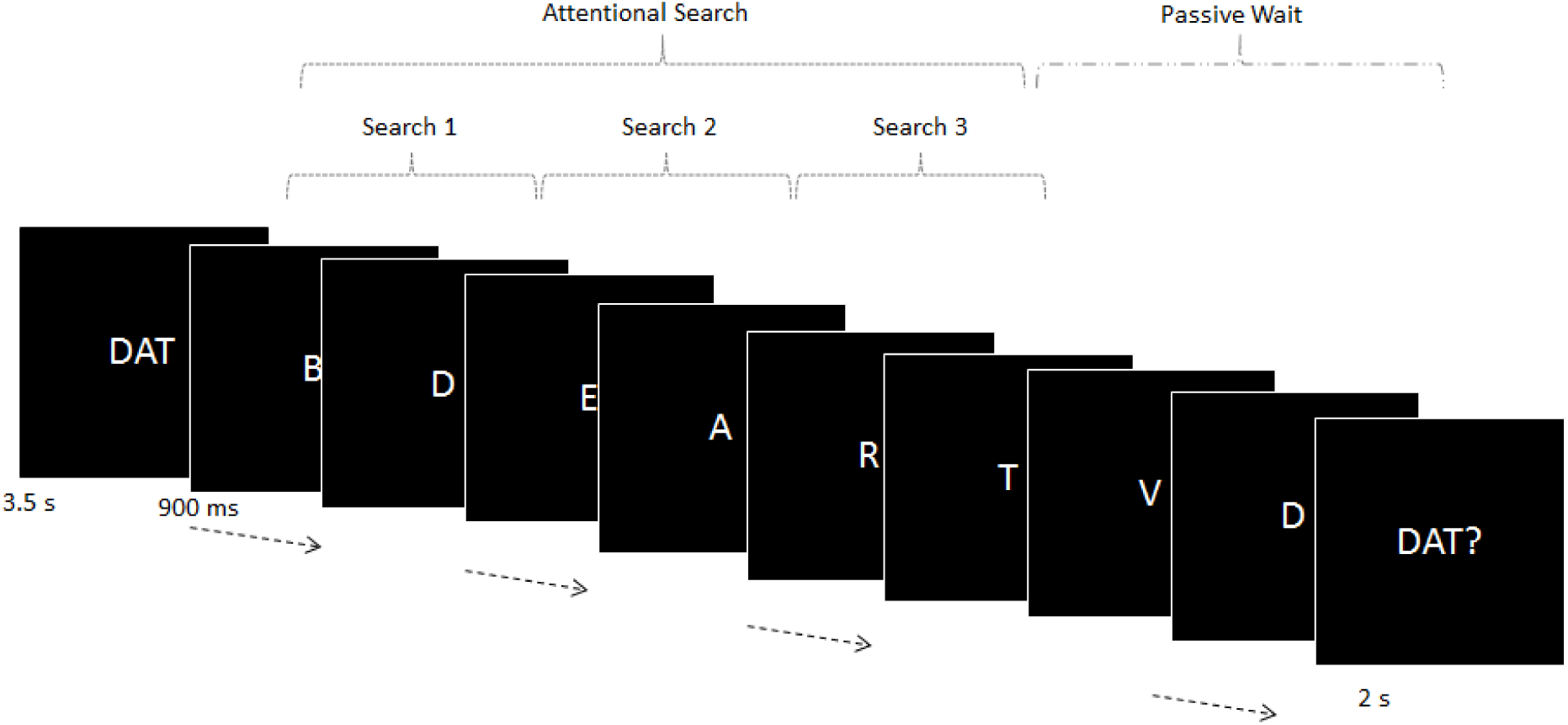
Experiment 1. Trials began with a three-letter cue word. The three letters of this word were to be covertly detected, in the correct order, in the ensuing letter stream. After all three had been detected search stopped and subjects waited for the letter stream to end, at which point a probe appeared asking if all three targets had appeared. In half of the trials all three targets did not appear and the search continued till the end of the sequence. The total length of the letter sequence was constant at 40 letters presented at the rate of 1/s (on for 900 ms). The length of the three searches and of the passive wait varied from 2 to 32 s.

Participants began by covertly monitoring for T1 (Figure 1, ‘D’), at its detection they looked for T2 and so on. The sequence of search was important; each target could only be searched for after the previous one had been detected and was irrelevant when it occurred before this point. After covertly detecting the three targets participants passively waited for the letter sequence to end. After which a probe appeared (e.g. ‘DAT?’) to which participants responded if all three targets had appeared in that trial. Responses were made on a button box positioned under the participant’s right hand (index finger for ‘yes’, middle finger for ‘no’). A variable inter-trial interval of 2—7 s preceded onset of the cue for the next trial. All three targets appeared in only 50% of the trials; in the rest one, two or none of the targets appeared. The inter-target interval within a trial varied randomly between 2 to 32 s.

All stimuli were centered on the screen, visible from the participant’s position in the scanner via a mirror mounted within the head coil. Letters were presented in white fonts on a black background and subtended a visual angle of 2° vertically. Participants learnt the task in a 10 min pre-scan practice session and then proceeded to a scanning session of an hour, which was divided into three separate scanning runs, each consisting of 20 trials.

We investigated if attended perception of the non-target letters of the sequence elicited fronto-parietal activity. In the first analysis we modeled the entire 40 s letter sequence with a set of 20 finite impulse regressors (FIR) starting from its onset. Target events were separately modeled with eight 2 s long FIRs. This allowed us to delineate the activity during the perception of the letters of the sequence as well as that elicited by the target letters without making assumptions about the nature of the canonical hemodynamic response.

In the second analysis we contrasted the activity elicited when letters were attentively perceived (i.e. during the attentional search) with that during the passive wait after the search was over (i.e. when letters were no longer under top-down attention). For this the periods of the three searches and the passive wait were modeled with separate epoch regressors, while target events were modeled as events of no duration. These regressors were convolved with hemodynamic basis function (HBF).

In both analyses, the cue and the probe were modeled using epoch regressors of width equal to their durations and convolved with the HBF. Movement parameters and block means were included as covariates of no interest. Parameter estimates for each regressor were calculated from the least-squares fit of the model to the data, and (in the second analysis) estimates for individual participants were entered into a random effects group analysis.

Whole-brain comparisons were performed using paired t-tests on the relevant contrast values from each participant’s first-level analysis. Unless otherwise specified, all results are reported at a threshold of p < 0.01 and corrected for multiple comparisons using the false discovery rate. Coordinates for peak activation are reported using an MNI template.

To capture frontoparietal regions widely engaged in cognitive control, 10 regions of interest (ROIs) were created as spheres of 10 mm radius at coordinates that have been shown to be consistently active in varied tasks (Dosenbach et al., 2006; Duncan, 2006). Note these are also the regions seen active in a number of previous studies on neural correlates of consciousness (bilateral IFS, IPS, AI, APFC; reviewed in Rees, 2007). The ROIs (in MNI space) were bilateral inferior frontal sulcus (IFS; central coordinate ± 41 23 29), bi-lateral intraparietal sulcus (IPS; ± 37 – 56 41), bilateral anterior insula extending into frontal operculum (AI/FO; ± 35 18 3), anterior cingulate cortex (ACC; 0 31 24), and presupplementary motor area (pre-SMA; 0 18 50), all taken from Duncan (2006); bilateral anterior prefrontal cortex ROIs (APFC; 27 50 23 and – 28 51 15) were taken from Dosenbach et al. (2006). ROIs were constructed using the MarsBaR toolbox for Statistical Parametric Mapping or SPM. Estimated data were averaged across voxels within each ROI using the MarsBaR toolbox, and the mean values were exported for analysis using SPSS.

The overall scheme of the experiment was similar to experiment 1 (Figure 2). Here, we compared the activity elicited during attended viewing of the letter sequence to passively waiting in front of a blank screen. At the beginning of each trial participants were shown a 3-digit sequence (e.g., ‘517’ presented for 1 s). They were to sequentially detect each of the three digits in the ensuing letter sequence. Between 1 and 7 seconds later (jittered, mean = 3 s) a blue square (2.5° visual angle) appeared on the screen which either remained blank till the appearance of the first target number (blank phase) or contained a series of consecutive letters (presented in black font on white background, visual angle of 2° vertically) each presented for 1 s with no intervening gap (letters phase). Numerical targets and letters were identical in font, color, size and duration. At the appearance of the 1st target (T1) participants pressed the button under their right index finger and started searching for T2. At its appearance they pressed their right middle finger button and began searching for T3, which appeared in 50% of trials in which case they pressed their ring finger button. The remaining trials ended without T3 being presented. Half of the search periods leading up to T1 and T3 (1st and 3rd phases) were blank during which participants passively waited for the target to appear, while the period between T1 and T2 (2nd search phase) was always a letter phase. Trial length was 30 s. The length of the 1st and the 3rd phase was jittered between 2 to 22 s while that of the 2nd phase varied between 2 to 7 s.

**Figure 2.**
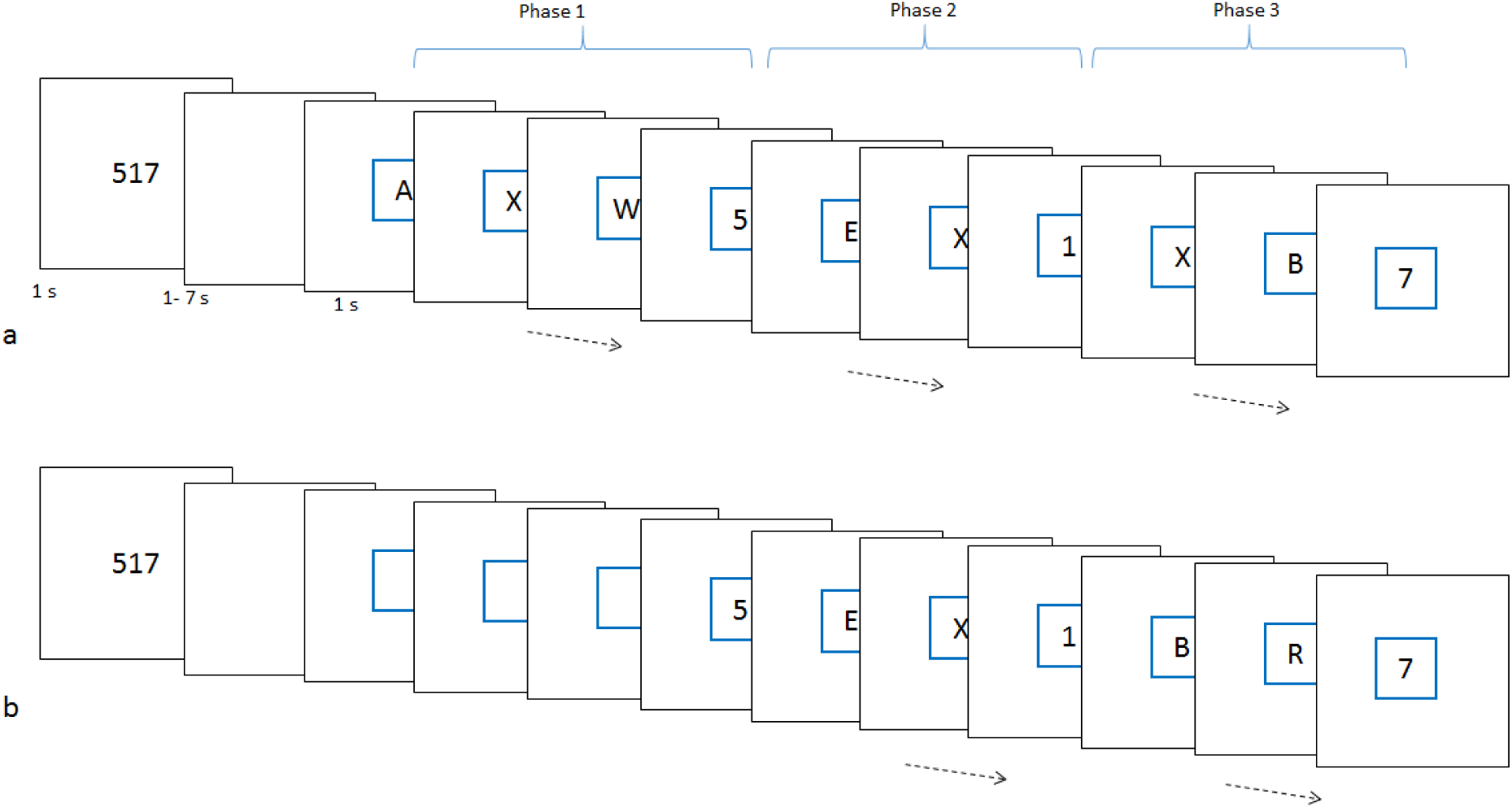
Experiment 2. Trials began with the presentation of a three digit number cue, appearances of which were to be sequentially detected in the following search episode. Search started after an average gap of 3 s and was marked by the appearance of a blue square, which either remained blank during blank phases or contained a series of letters (letter phases) each appearing for 1 s with no gap between consecutive letters. Targets were presented in identical conditions to the letters. Participants made button press responses on detecting the correct target. Note that in the above figure the third (a) and first (b) phases are blank.

Participants were told that correct and fast responses to all 3 targets (when present) would score 1 point. Responses to only two targets when only two were presented would not score. Any trial which contained a response to a non-target or a non-response to a target would lose a point. The current score was displayed at the end of each trial for 1 s. The next trial began after a variable inter-trial interval of 2 - 7 s.

Letters were selected in a quasi-random manner such that a particular letter (‘X’ or ‘Y’) was presented with a higher probability (0.2). This allowed a check at the end of the session as to whether participants had noticed one letter appearing more frequently than others without giving prior warning that any letter report would be required (see below).

We did two kinds of analysis. In the first Letter and blank phases were modeled with epoch regressors while target, cues and score events were modeled as event regressors. These were convolved with the HBF. Movement parameters were included as covariates of no interest. Parameter estimates for each regressor were calculated from the least-squares fit of the model to the data, and estimates for individual participants were entered into a random effects group analysis.

In the second analysis twenty seconds of activity following the beginning of first phase was modelled with ten 2s long FIR regressors.Letter phase and blank periods were separately modelled. Targets and cue were modelled as events of no duration and convolved with the HBF. Through this we aimed to capture the activity elicited during the first phase. We chose this period over the third phase because activity captured by FIR model during the third phase will inevitably also include that elicited by the third target detection.

### Experiment 3

In this experiment we used a very different design. The perceptual events investigated in the previous two experiments were clearly visible and had to be attended to but their presence did not have much consequence for the subsequent course of cognition. In this experiment we investigated if cues that informed the rule to be used to achieve the goal elicited fronto-parietal activity.

Subjects waited for one of the four auditory cues (‘P1’, ‘P2’, ‘F1’ and ‘F2’ spoken in a mechanical voice) while viewing a fixation cross (Figure 3). The cue informed which of the four rules were to be used for responding to the visual stimuli to follow. After a variable wait of 1.5 to 9 s the visual stimuli appeared. They consisted of a picture of a face and of a place placed above/below the fixation cross. Depending on the cue participants categorized the following stimuli. P1 - the picture of place as indoor (index finger) or outdoor (middle finger); P2 - place as indoor commercial (index), indoor non-commercial (middle), outdoor commercial (ring) and outdoor non-commercial (little finger); F1 - face as male (index) or female (middle finger); F2 - face as young male (index), old male (middle), young female (ring), old female (little). The loudness of the cue was set at a level that was comfortably and clearly audible.

**Figure 3.**
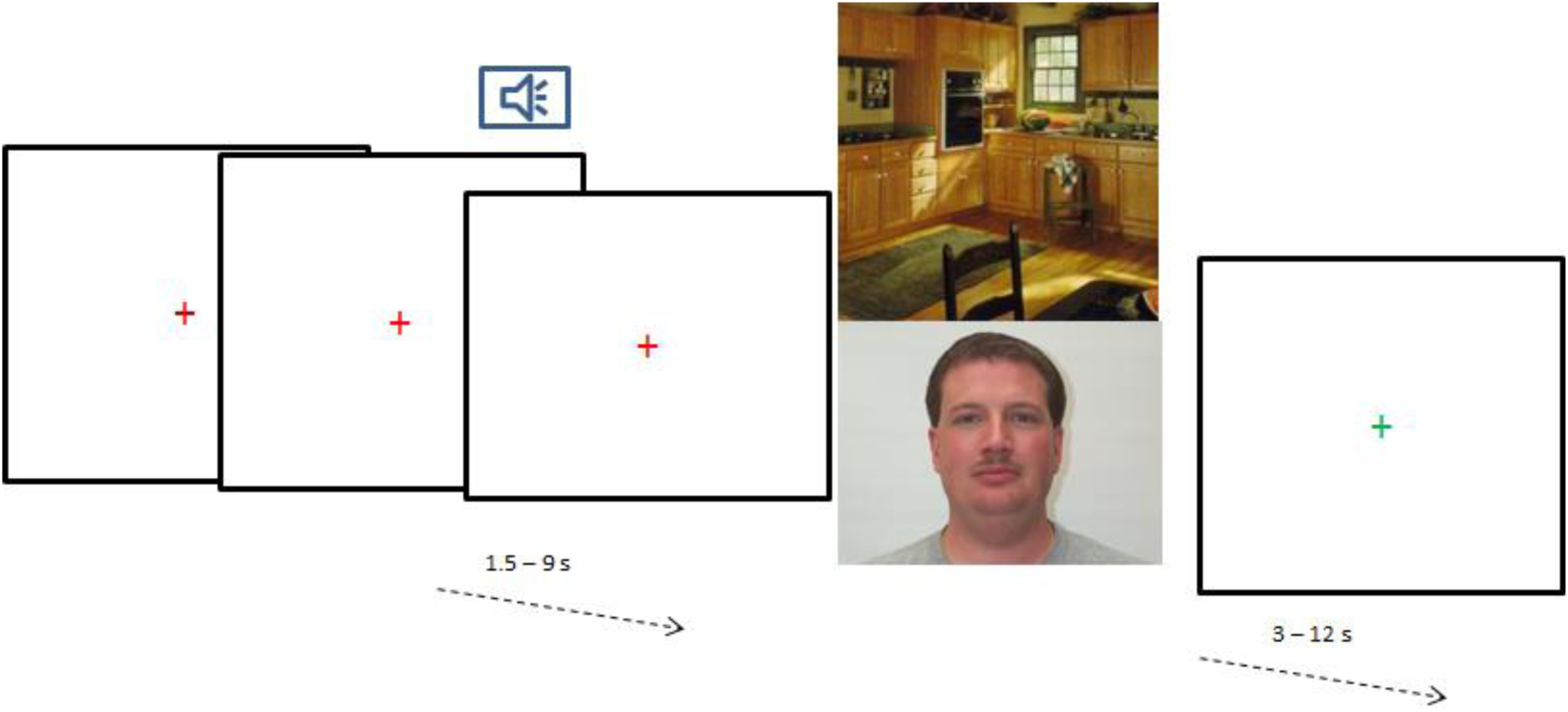
Experiment 3. Trials began with an auditory cue that informed the rule to be used in categorizing the relevant picture in the visual stimuli to follow (see text). Four potential rules were cued by four different cues. After the response to these stimuli fixation-cross color changed signaling the beginning of inter-trial period. In 20% of trials the visual stimuli did not appear, and the auditory cue and the wait for the visual stimuli were directly followed by a change in color of the fixation cross signaling the end of the trial.

All responses were made with right hand. After the response the color of the fixation cross changed signaling the end of the trial. The next cue followed a random interval of 3 to 12 s. On one-fifth of the trials the cue was not followed by the stimuli instead after a random wait the color of the fixation cross changed without the stimuli appearing, and the inter trial trial began with subjects waiting for the new cue to appear. This was done to further separate the activities elicited by the cue from those related to the stimuli.

We investigated the time series of activity elicited by these cues that were clearly perceived, consciously attended and task relevant; in fact their information was to be maintained for up to next 9 s. The cue was clearly not the goal of the task, but nonetheless was indispensable for goal completion. We modeled the period of 10 s following the cue with ten 1 s long FIRs. Unlike the cue, the response coincided with the completion of the task goal. We likewise modeled the 10 s period following the response with ten FIRs. Rest of analysis was similar to experiment 1.

### Subjects

Twenty-one participants were recruited for each of experiments 1 (15 females; mean age, 24.5 ± 4.1years) and 2 (12 females; mean age, 22 ± 4 years) and twenty seven participants were recruited for experiment 3 (15 females; mean age 50 ± 6.6 years). In experiment 3, two participants had to be excluded from analysis due to very poor behavioral performance. However, including their data did not change the results from that reported here. All participants were recruited from the MRC-CBU volunteer panel. Participants were right handed and had normal or corrected vision. Informed consent was taken, and the participants were reimbursed for their time. The study had the approval of Hertfordshire Local Research Ethics Committee.

### Acquisition (Experiments 1 and 2)

fMRI data were acquired using a Siemens 3T Tim Trio scanner with a 12 channel head coil. A sequential descending T2*- weighted echo planar imaging (EPI) acquisition sequence was used with the following parameters: acquisition time, 2000 ms; echo time, 30 ms; 32 oblique slices with slice thickness of 3 mm and a 0.75 mm interslice gap; in-plane resolution, 3.0 _ 3.0 mm; matrix, 64 _ 64; field of view, 192 mm; flip angle, 78°. T1-weighted MP RAGE structural images were also acquired for all participants (slice thickness, 1.0 mm; resolution, 1.0 _1.0_ 1.5 mm; field of view, 256 mm; 160 slices). Experimental task started after 12 “dummy” scans had been acquired. These were discarded from the general linear model to allow for T1 equilibration effects.

Experiment 3 – Acquisition was identical to the above except that 16 slices were taken in an acquisition time of 1000 ms. Acquisition window was angled to include occipital, parts of temporal (specifically auditory regions) and frontal regions leaving out superior parts of frontal and parietal cortices, anterior temporal regions and inferior cerebellum.

### Analysis

The fMRI data were analyzed using SPM8 and SPM12 (experiment 3). Before statistical analysis, all EPI volumes were slice-time corrected using the first slice as a reference and then realigned into a standard orientation using the first volume as a reference. These realigned volumes were then normalized into the Montreal Neurological Institute (MNI) space and spatially smoothed using an 8 mm full-width half-maximum Gaussian kernel. During the normalization stage, voxels were resampled to a size of 3_3_3 mm. The time course of each voxel was high pass filtered with a cutoff period of 90 s.

## Results

### Experiment 1

Average response time was 770 ± 18 ms, while average accuracy was 96.1% (±2.7). Expectedly, target detection events elicited strong activity in all fronto-parietal regions, with the final target eliciting higher and more widespread activity (colored plots in Figure 4; see Farooqui et al. 2012 for a detailed analysis). But our concern in this study was the activity elicited by the non-target letters. Such letters were consciously perceived - they were presented at fixation for long duration and had to be attended to detect target letters in their midst. If attended and conscious perception of such letters elicited increased fronto-parietal activity we would expect increased activity during the visual search than during the preceding or succeeding periods of rest.

**Figure 4.**
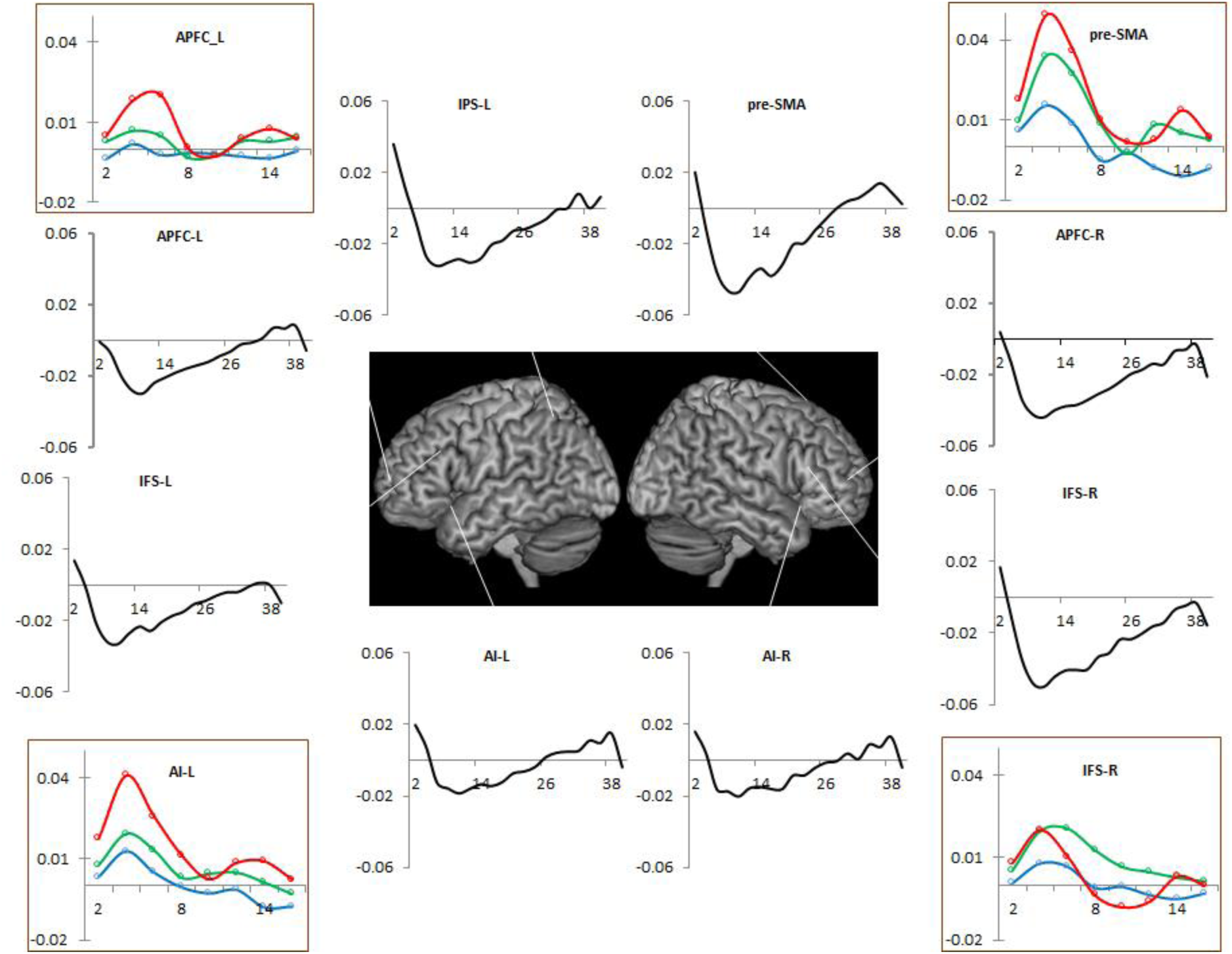
Experiment 1. Black plots show the estimated activity during the 40 s of letter sequence presentations. Note that the start of the sequence was accompanied by widespread deactivation that started returning towards the baseline after 12 - 14 s. In contrast targets elicited strong activity (red: T3, green: T2, blue: T1).

Figure 4 shows the time series of activity starting from the beginning of the letter stream. None of the fronto-parietal regions increased their activity with the onset of the letter stream. Instead all of these regions deactivated and continued to do so for up to initial 14 s and then returned to the baseline towards the end of the search. Sequential conscious perception of the letters was not correlated with an increase in the activity of these regions. In fact even an event as salient and conscious as the beginning of visual search did not activate these regions, instead coincided with a widespread deactivation.

We next compared activity during the period when the letter sequence was being attended to (i.e. during the search) to when they were passively viewed (i.e. after the search had ended). Only visual regions near the occipital pole showed greater activity during attentive search compared to the subsequent passive wait (Figure 5). In contrast, very widespread set of regions that included both cognitive control and default mode related fronto-parietal regions showed greater activity during passive wait compared to the attended search.

**Figure 5.**
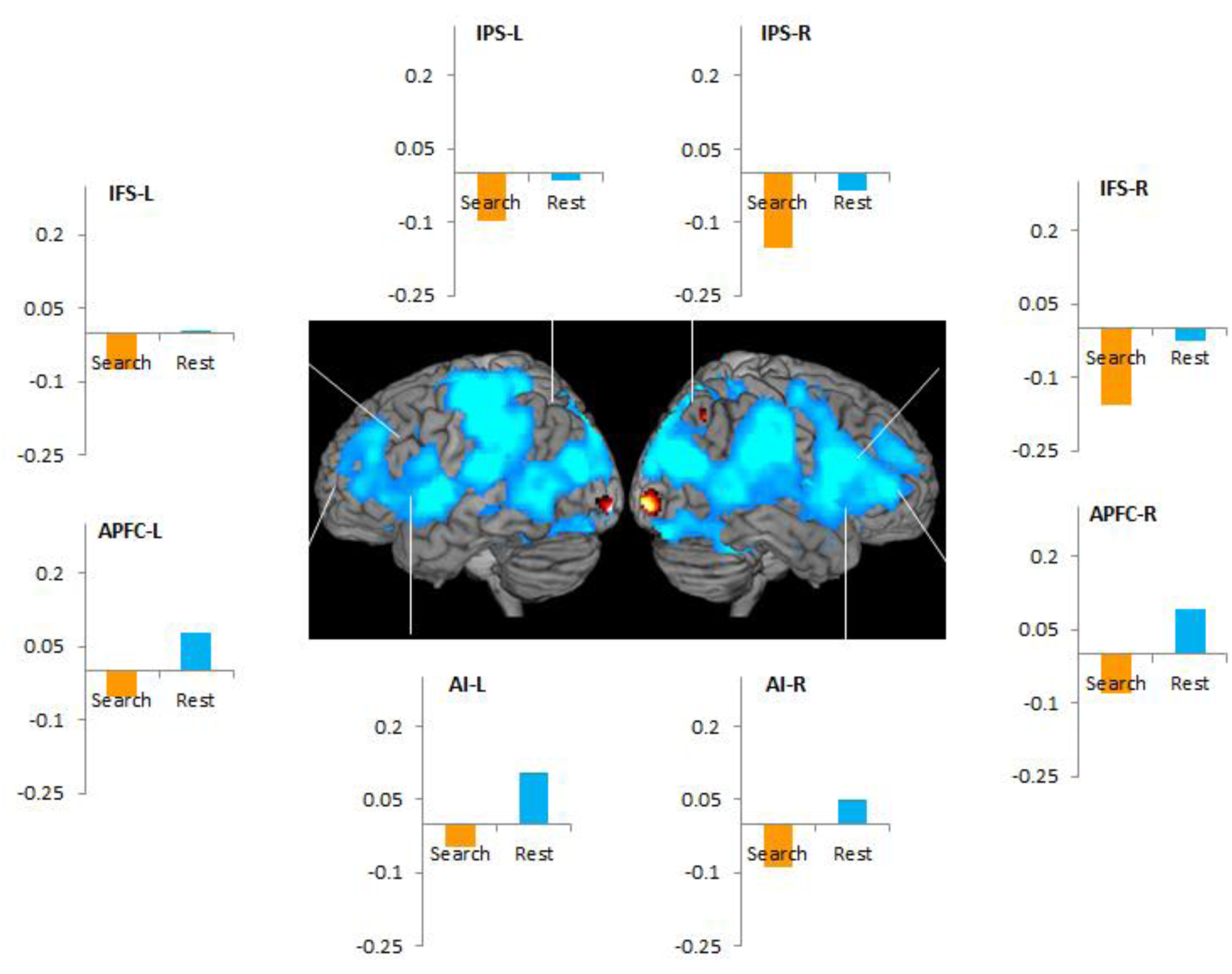
Experiment 1.‘Cooler’ regions in the whole brain render were more active during rest following the search than during search, while ‘hotter’ regions were more active during the search. Plots show that all fronto-parietal regions were more active during the rest period than during the search.

Thus, the conscious and attended perception of the letter events correlated with a decrease in fronto-parietal activity compared to both the preceding and succeeding intervals when subjects were waiting passively. Through the next experiment we show that the fronto-parietal activity elicited during the conscious and attended perception of these letters was lower than during waiting passively in front of a blank screen.

### Experiment 2

Most participants exceeded 97% (average 98.2 ± 0.3) with a mean reaction time of 410 ± 10 ms. RT of detecting targets in the midst of the letter sequence was faster than when targets appeared after blank waiting periods (t_20_ = 2.2; p = 0.03). This would be expected because during the letter sequence participants were actively engaged in search and were ready for the possibility that any letter event could be a target, while during blank periods they were passively waiting for the target to appear and can be expected to be prone to mind wandering (Smallwood et al. 2008). Post experiment 17/21 participants could correctly recall the letter that appeared most frequently in the letter sequence. The remaining 4 who were not certain of their answer, nonetheless, answered correctly when given four options to choose from.

Again, if increased fronto-parietal activity is integral to attended and conscious perception then we would expect higher activity in these regions during letter sequence phases when participants are attending and perceiving a series of letters than during blank periods, during which the perceptual phenomenon was passive and constant. However, a whole brain contrast (Letter phase > Blank phase) was significant only in occipital regions (Figure 6), wherein it extended from the occipital pole (BA 17) laterally through BA 18, 19 and 37 (the lateral occipital complex, LOC; Grill-Spector et al. 2001). Small clusters of significance were seen subcortically in head of caudate and posterior thalamus. Decreasing the statistical threshold (to uncorrected p <0.05) showed an additional small cluster in left posterior prefrontal region.

**Figure 6.**
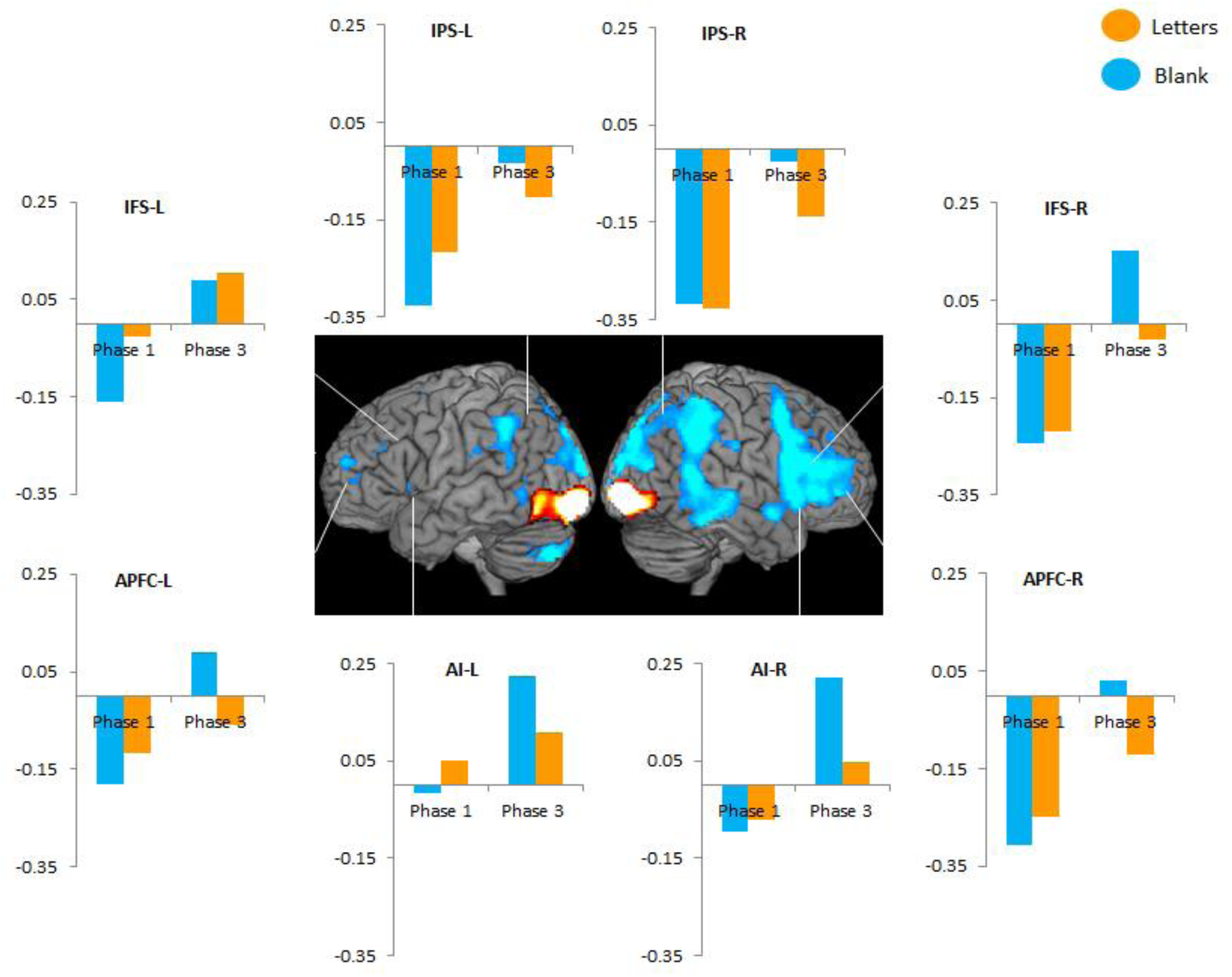
Experiment 2. Hotter and cooler regions were more and less active respectively during the search through the letter sequence than during the passive wait in front of the blank screen. ROI analysis showed that during phase 1 only left IFS showed greater activity during letter sequence than blank phases, other ROIs did not differ. However, during phase 3 most ROIs showed greater activity during blank phases compared to the letter sequence.

A reverse of this contrast (Figure 6) showed that activity during the Blank phases was greater than the Letter Phases in widespread brain regions including right fronto-parietal cortices, medial occipital and parietal regions (Calcarine sulcus, lingual gyrus, cuneus and precuneus), temporo-parietal junction, inferior parietal lobule, again suggesting that the attended and conscious perception of letters were accompanied by widespread deactivation of fronto-parietal regions.

Plots in Figure 6 show the activity elicited in these regions by the Blank and Letter sequence phases when they corresponded to the first and the third phases. During the 1st phase, almost all regions showed a net deactivation. In most ROIs the activity elicited during the Letter phases was not different from Blank phases. Only in left IFS (paired t_20_ = 2.3, p = 0.03), with some trends in left IPS (p = 0.07) was the activity during Letter phases significantly less negative than during Blank phases. It is noteworthy that none of these regions showed net positive change in activity (see below for further analysis on this issue).

All regions elicited higher activity during the third phase compared to the first phase (see Farooqui & Manly, under review, for a detailed analysis of this issue). Importantly, the Letter sequences during the third phase elicited less activity than the Blank phases of this search in all ROIs except left IFS. This reached statistical significance in left AI and APFC, right APFC, IFS, IPS and AI (t_20_ > 3.7, p<0.01). Importantly in left IFS there was no difference between these two phases (t_20_ = 0.3, p = 0.7).

Through a separate GLM, we looked at the time course of activity starting from the beginning of the search through next 20 s (corresponding largely to the first search period) by modeling them with ten 2-second long FIR regressors. Figure 7 confirms that in the fronto-parietal ROIs including the left IFS both Blank and Letter Phases were linked to a deactivation starting with the onset of search and the presentation of the letter events.

**Figure 7.**
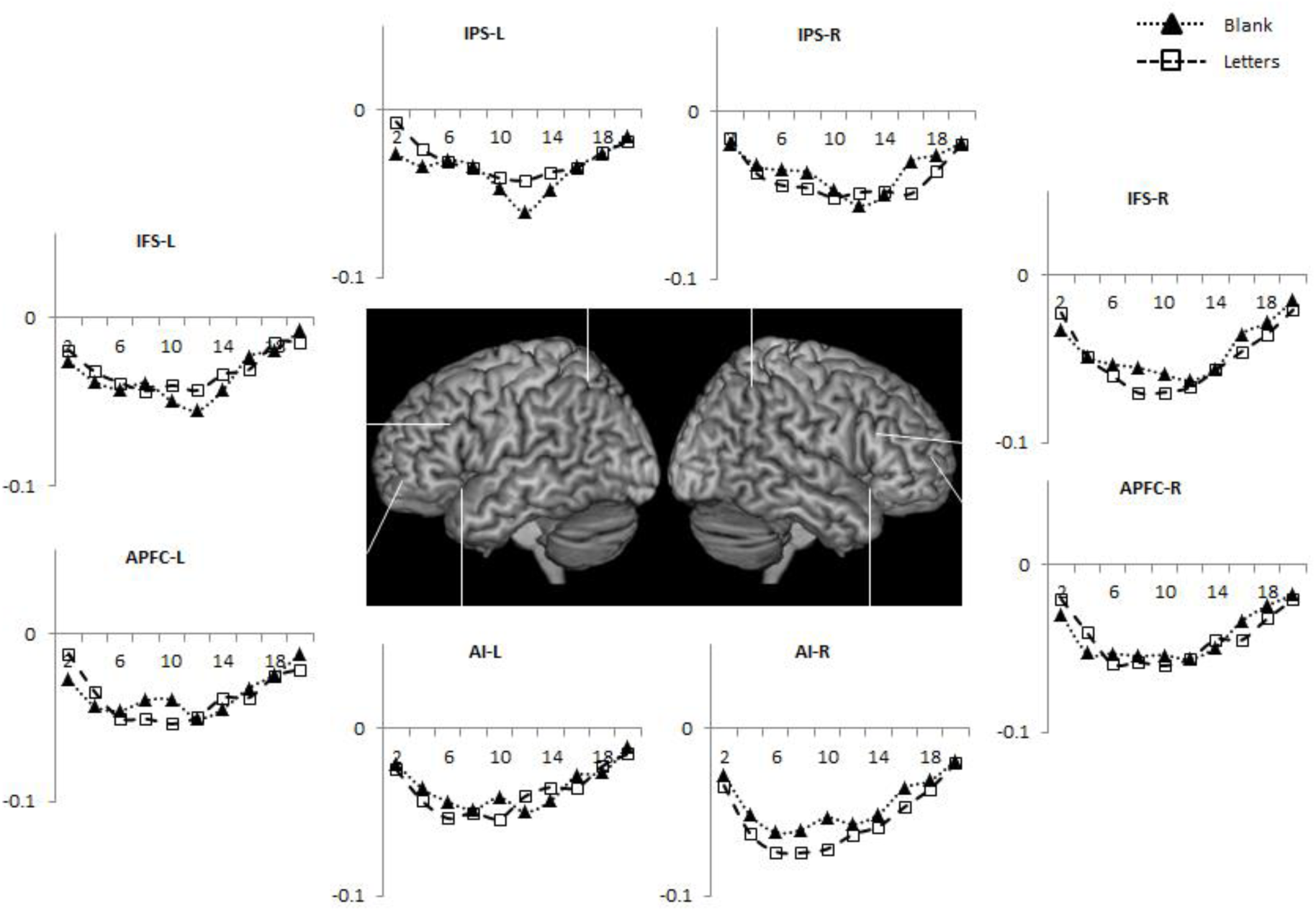
Experiment 2. As in experiment 1 early part of the search was accompanied by a widespread deactivation of fronto-parietal regions that gradually returned towards the baseline.

Again, attended and conscious perceptual events (the letters of the sequence) that were not goals of ongoing tasks instead of activating were accompanied by a deactivation of fronto-parietal regions. This was self-evident during the first phase and could be inferred during the third phase because the absence of these letters elicited greater fronto-parietal activity.

### Experiment 3

Perceptual events in experiments 1 and 2 though attended and consciously perceived were nonetheless inconsequential for subsequent cognition because the non-target letter event did not affect the subsequent course of task execution. In experiment 3 we show that the perception of cues that have to be kept in working memory for extended periods of time and that inform the rule to be used for subsequent goal completion does not elicit increased fronto-parietal activity either.

Average accuracy and RT were 89.3 % (± 1.1) and 2660 ms (± 241). Maroon plots in Figure 8 show the estimates of activity following the cue. As is evident, increased activity was seen only in auditory regions (A1) and not in any prefrontal region. In contrast, the response to the stimuli–marking the completion of the goal – elicited widespread prefrontal activity (blue dashed plots).

**Figure 8.**
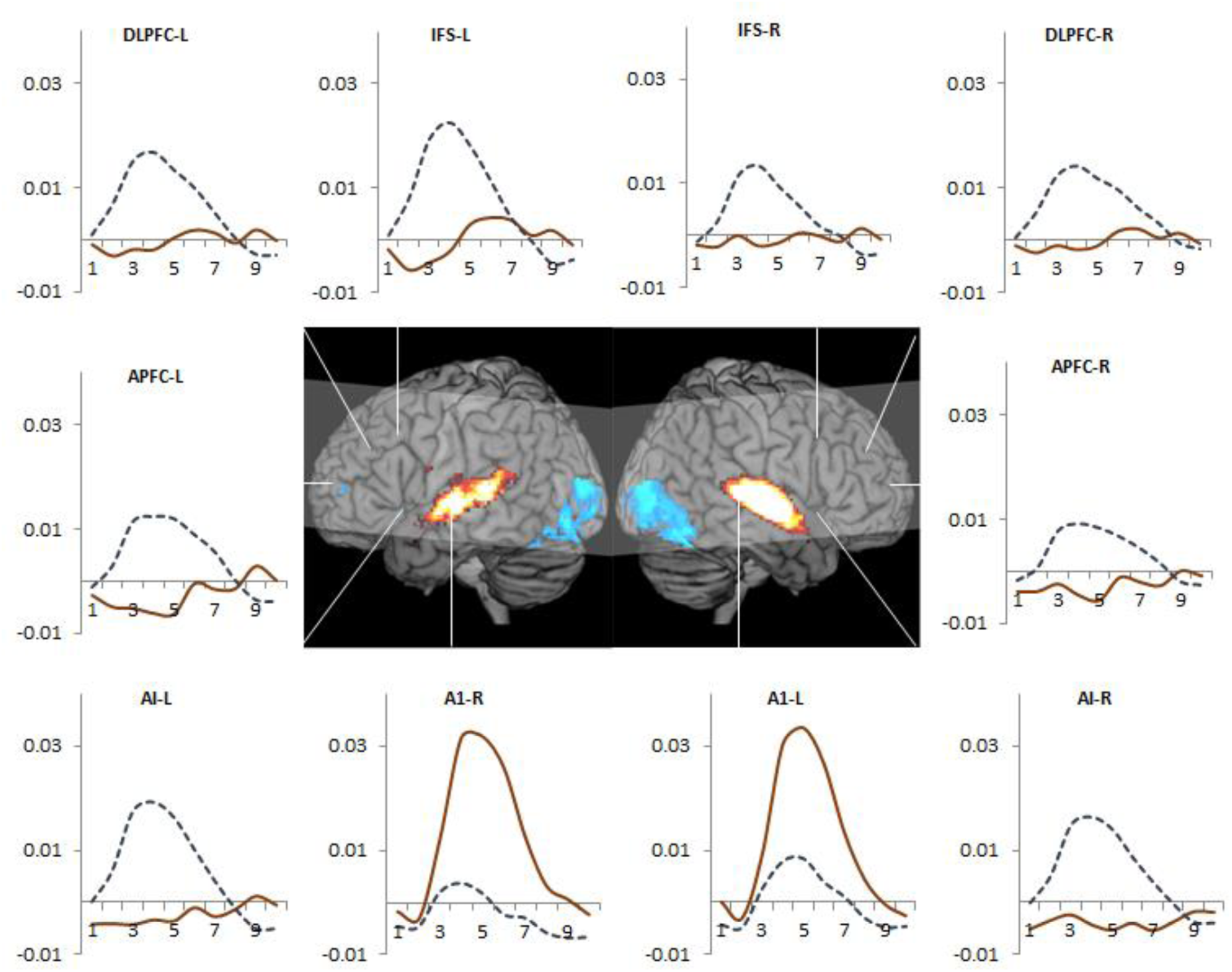
Experiment 3. Maroon plots show the average estimate of activity elicited by the auditory cues. Note that these cues only activated the auditory regions. In contrast, the response event (dashed blue plots) activated all fronto-parietal regions. In brain render the hot and cold colors show regions where the BOLD activity increased and decreased across initial 5 seconds following cue perception.

Next we contrasted activity estimates for each of the scanned voxel during the first two seconds to that during the fourth and fifth seconds following the cue. This would capture any scanned region where perception of cue elicited an increase in activity. However, this analysis too yielded significant results only in auditory temporal regions (brain render in Figure 8; results shown at a liberal threshold of uncorrected p <0.01). Thus, prefrontal activity did not increase even in response to highly salient cues that had to be remembered for extended intervals. Note that unlike experiments 1 and 2, attended cues did not deactivate fronto-parietal regions.

## Discussion

Previous research has linked conscious and attended perception to increased front-parietal activity (Duncan, 2006; Rees, 2007; Dehaene and Changeux, 2011). Here, such perception delinked from task goals elicited a widespread deactivation of fronto-parietal regions (experiments 1 and 2) or showed an absence of increased activity (experiment 3). While arguing against the necessity of increased fronto-parietal activity for conscious and attended perception, current results suggested that goals may have been the key confounds in the purported association between them.

### Consciousness

Though our experiments could not specify regions whose activity correlated only with the conscious perception, they decisively tested the thesis that fronto-parietal activity is a necessary correlate of conscious perception. Our negative results suggested that previous results were indeed confounded by issues related to resolving perceptual ambiguity, goal completion, and explicit and difficult metacognitive judgments. This is also supported by an analysis of previous studies that did not find fronto-parietal activity during conscious perception.

Tse et al. (2005) found that conscious perception of unattended and task irrelevant stimuli correlated with visual regions and not with fronto-parietal activity. Like the current study this study did not ask participants to report their awareness and relied on perceptual conditions where conscious perception was extremely likely. This may be important because the requirement to report phenomenal awareness makes detecting changes in awareness a goal akin to detecting targets during a search. Participants have to explicitly monitor their awareness for changes linked to the perception of target event e.g. shift in current percept during binocular rivalry and when that occurs their goal of the metacognitive task is complete - they have detected what was required to be detected. Consequently, experiments that did not involve explicit phenomenal reports found decreased to absent prefrontal activity during conscious perception (Tsuchiya et al. 2015). For example, Frassle et al. (2014) showed that prefrontal activity seen during binocular rivalry decreased when the requirement to report the current percept was removed, suggesting that issues related to metacognitive monitoring, reporting the subjective state, and related action selection at least partially accounted for such activity.

Many studies have relied on degraded or difficult to interpret stimuli e.g. during visual masking and binocular rivalry, and the accompanying fronto-parietal activity may have been related to the success in resolving perceptual difficulty. This was evident when Moustoussis and Zeki (2002) used perceptual ambiguity to render stimuli unconscious. Pictures made through color contrasts could either be easily visible e.g. green house on a red background, or they were rendered non-conscious through binocular rivalrous presentation of identical stimuli made with opposite color contrasts (green house on red background in one eye and red house on green background in the other). They found that easily visible pictures activated only visual regions while the ambiguous non-conscious pictures activated prefrontal regions. Akin to the current results, Goldberg et al. (2006) showed that regions that activate during explicit introspection (which included many fronto-parietal regions) deactivate during simple perceptual categorization that was likely to involve conscious perception.

It is possible that cortical regions that process the perceptual and task related aspects of a stimulus may also be the ones that make it conscious. Identifying difficult to interpret stimuli (e.g. during visual masking and binocular rivalry) is difficult and requires widespread neural processing that also correlate with their conscious perception. In contrast, letters in experiments 1 and 2 were easy to process and inconsequential for goal directed action and hence activated only early regions of visual processing hierarchy. This thesis was well demonstrated when participants were asked to identify pictures of animate/inanimate objects made with fragmented colored lines and hidden amongst other lines (Eriksson et al. 2008). Consciously identifying was initially difficult and activated widespread fronto-parietal regions, but as identification became easier with practice and this activity reduced. Likewise, when pictures of fearful faces were presented in binocular rivalry with disgusted or neutral faces, they were more likely to win the competition and be the dominant conscious percept but, crucially, elicited less fronto-parietal activity than other face pictures (Amting et al. 2010). Case reports of patients with bilateral frontal lobotomy suggest that these patients did not lose their conscious perception (Miller, 1967; Fleming and Dully, 2008), nonetheless, patients with frontal damage do show a deficit in perceiving degraded and fleeting stimuli (Del Cul et al. 2009).

### Attention

Were these events attended? While it would be difficult to argue that the cues in experiment 3 were unattended, it may be claimed that the letter events in experiments 1 and 3 were not really attended. Even if this claim was accepted for a moment, it is still notable that the *onset* of the letter sequence – an event very likely to be both salient and attended (like the cue in experiment 3) - also elicited a fronto-parietal deactivation.

The construct of attention is used for task events (space and objects) that are selectively focused upon in a top-down manner to the exclusion of task irrelevant events (James, 1890; Desimone & Duncan, 1995). Here the letters of the sequence were task *relevant* in that they were part of the relevant task environment and were relevant task events as opposed to ongoing task irrelevant peripheral vision, scanner noise, various somatic sensations etc. Most importantly, the current results showed that these letter events were not just ignored as unattended but elicited a deactivation of fronto-parietal regions.

Note that the current results need not imply that these regions were not needed for attentional functions of identifying and rejecting non-targets (experiments 1 and 2). It remains plausible that these regions were needed to create attentional task sets early on in the experiment, which then filtered out non-targets early in visual processing without the need for an ongoing increase in fronto-parietal activity (for related ideas see Hommel, 2000; Muhle-Karbe et al. 2016). Such sets could have been maintained across time without sustained increased fronto-parietal activity (e.g. Lewis-Peacock et al. 2012; Stokes et al. 2013), perhaps through a sustained change in synaptic weights within the relevant network (Mongillo et al. 2008). The decrease in activity could have been a means of preserving synaptic weight configurations maintaining the attentional set by preventing irrelevant activity. Alternately, deactivation could represent active exclusion of inconsequential task events from fronto-parietal cortex because of limited representational capacity of these regions (see Marois & Ivanoff, 2005) and the potential of fronto-parietal representations to influence processing in widespread regions (Duncan, 2006).

Attention is not just an amplifier of sensory representation being attended to, and may be better seen as a means of ensuring that incoming percepts are assigned to neural systems where their subsequent processing will enhance goal completion, and that they are excluded from systems where their representation/processing is not required or may be disruptive. Hence, salient and attention grabbing distractors are demonstrated to be better excluded from deeper cognitive processing than less salient distractors (Hickey et al. 2009; Gasper et al 2014; Moher et al. 2015; Gaspelin et al. 2015). While these studies demonstrate the active exclusion of attention grabbing events that appear at the same time but at a different space from the targets of the search or goal of the action, current results suggested the possibility of suppression mechanisms that operate across time while the target is being searched for.

This view of attention may also explain frequent observations of deeper processing of task irrelevant events that are less visible or subliminal and better exclusion of those that are more visible. Thus, the learning of associations involving subliminal and incidental perceptual events in the task environment is better than associations involving more visible events (Watanabe et al. 2001; Tsushima et al. 2006, 2008; Farooqui & Manly, 2015). Likewise, subliminal events presented during the execution of an ongoing task frequently results in stronger priming effects than more visible events (Lau & Passingham, 2007; Zhou & Davis, 2012; Manly et al. 2014; Lin & Murray, 2015). Arguably, in such cases more visible events are more accessible to attentional mechanisms that put stronger limits on their processing compared to the case with subliminal events.

Current study showed that conscious and attended perception far from necessarily activating fronto-parietal regions can be accompanied by a deactivation of these regions when delinked from task goals. This raised a speculative possibility of an attentional suppression mechanism that excludes inconsequential task events from achieving widespread processing.

## Acknowledgments

This work was supported by United Kingdom Medical Research Council Grant MC-A060-5PQ20.

